# *Odd-skipped* controls neurite morphology and affect cell survival in *Drosophila Melanogaster CNS*

**DOI:** 10.1101/2020.02.11.943373

**Authors:** Yeoh Sue Lynn, Alina Letzel, Clemence Bernard Hannah Somerfield, Kyle Kyser, Emily Lin, Amanda Roper, Yucen Yuan, Chloe Saunders, Mina Farag, Samual Colourous, Camilla W. Larsen

## Abstract

The transcription factor *Odd-skipped* has been implicated in many developmental processes in *Drosophila melanogaster*. *Odd-skipped* is expressed in a small cluster of neurons (Slater, Levy et al.) in the developing and adult CNS but its role in neurogenesis has so far not been addressed. Here we show that *Odd-skipped* plays a pivotal role in neurite growth and arborization during development. Loss-of-*Odd-skipped* function prevents neurite outgrowth whereas over and miss-expression causes neurite growth and arborization defects. In addition, miss-expression of *Odd-skipped* can induce cell death in some neural sub types. The neurite growth and arborization defects associated with *Odd-skipped* over expression correlates with a reduction in the pre-synaptically targeted protein Bruchpilot in axonal arbours suggesting an overall decrease in Odd neural synapse formation. This is supported by behavioural data showing that larvae in which *Odd-skipped* is overexpressed behave similarly to larvae in which Odd neurons are silenced showing that increasing *Odd-skipped* protein levels affect neural function. Finally, we demonstrate that using RNAi against *Odd-skipped* does not knock down *Odd-skipped* protein but instead cause an increase in protein levels compared to control larvae. This data demonstrates that RNAi can cause up-regulation of protein levels highlighting the importance of verifying protein levels when using RNAi approaches for knock-down.

## Introduction

Neurite development and growth requires a complex interaction between external factors, intrinsic properties and intracellular signalling and has been a subject of intense study both in vertebrates and invertebrates. In *Drosophila* for example, several different classes of genes including transcription factors, cytoskeletal proteins and guidance factors have been associated with dendritic and axonal growth and branching (Prokop, Beaven et al. 2013, Wang, Sterne et al. 2014). However, little is known about the molecular mechanisms underlying the first steps of neurite outgrowth in particular in the central brain. *Drosophila* is a particular well-suited organism to study the molecular machinery underlying neurite growth and branching due to the range of genetic tools which allow for detailed dissection of molecular interactions in defined subsets of cell. The *Drosophila* CNS develops in two phases; primary neurons are generated during embryogenesis and secondary neurons during larval and pupal stages. Primary neurons form the larval CNS whereas the adult CNS is composed by primary and secondary neurons. Although secondary neurons are generated during larval stages, they do not elaborate axonal and dendritic connection until metamorphosis and the larval functional CNS is therefore exclusively composed of primary neurons (Hartenstein, Omoto et al. 2020).

*Odd-skipped* (Odd) (Nusslein-Volhard and Wieschaus 1980) is a transcription factor which is part of the pair rule group of genes. Odd is expressed in a variety of tissues during *Drosophila* development (Coulter, Swaykus et al. 1990) and has been shown to play a role in the development of a range of different tissues. For example, Odd regulates epidermal segmentation, leg, eye and renal development (Coulter and Wieschaus 1988, Hao, Green et al. 2003Tena, 2007 #21, Bras-Pereira, Bessa et al. 2006) as well as maintaining prohemocyte potency (Gao, Wu et al. 2011). Similarly, the mammalian homologs *odd-skipped related 1* and *2* (OSR1 and OSR2) regulate urogenital, heart and bone development (Wang, Lan et al. 2005, James, Kamei et al. 2006, Kawai, Yamauchi et al. 2007). In addition, studies have shown that OSR1 act as a tumour suppressor gene in lung, gastric and renal cell carcinomas. OSR1 gene is downregulated in primary cancer cells as well as cancer cell lines and overexpression of OSR1 in gastric and lung cancer cell lines inhibits proliferation (Otani, Dong et al. 2014, Zhang, Yuan et al. 2017, Wang, Lei et al. 2018).

In *Drosophila* Odd is also expressed in the developing CNS. At embryo stage Odd is expressed in 2 of the Mushroom body MB (Levy and Larsen 2013) neuroblasts lineages as well as the CPM lineage. Odd expression in the MB is downregulated around the end of embryogenesis but is maintained in the CPM lineage which gives rise to the larval and adult Odd neurons (Levy and Larsen 2013, Slater, Levy et al. 2015). In early larval stages, Odd-skipped expression is restricted to 8 neurons although additional Odd positive neurons are generated in the lineage during larval and pupal stages. In the larvae 3 of the 8 Odd neurons project dendrites to the dendritic portion of the MB, the calyx, and all 8 neurons project axons to the CPM compartment situated ventral-medially in the brain (Slater, Levy et al. 2015). In the adult the Odd neural cluster project into various regions of the central brain including the calyx of the MB. In addition, 3 of the Odd neurons project into the lobular plate of the optic lobe (Levy and Larsen 2013) and these neurons respond to visual motion cues (Wasserman, Aptekar et al. 2015).

Here we show that Odd plays an important role in neurite growth and branching. Odd-loss of function prevents neurite outgrowth whereas Odd over and miss-expression affects neurite branching and arborisation and is associated with a decrease in the localisation of the pre-synaptic protein Bruchpilot at axonal terminals. In addition, Odd miss-expression affects cell survival since some neural sub populations are missing when Odd is miss expressed suggesting that Odd plays a role in cell proliferation and survival similar to the vertebrate homolog OSR1. Finally, we demonstrate that in the case of Odd-skipped, RNAi against Odd upregulates Odd protein level demonstrating that RNAi does not always knock down protein levels.

## Materials and Methods

### Fly strains and genetics

All *Drosophila melanogaster* fly strains used in this study were maintained at 25 °C on standard fly food and a 12-hour day/night schedule. Unless otherwise stated all transgenic lines were obtained from Bloomington Drosophila Stock Center (BDSC), Indiana University, IN. An Odd-Gal4 line (kind gift of Fernando Casares) (Bras-Pereira et al., 2006) was used to drive UAS-RNAi VDRC (Vienna Drosophila Resource Centre ID 107257) and UAS-RNAi (TRiP) 28295 and 34328 (Perkins, Holderbaum et al. 2015). Fly crosses for all RNAi and over expression studies were maintained at 29 °C as well as the control Odd-Gal4 fly strain. Unless otherwise stated UAS-CD8::GFP was included in all crosses to allow for morphological characterisation of neurite arbours. For the VDRC RNAi line and the TRiP RNAi line 28295 UAS-Dicer 2 was included in the cross. Control MARCM clones (Lee and Luo 2001) were generated using the following stocks: FRT19A, Odd-Gal4, UAS-CD8GFP and FRT19A, hsFLP, Tub-Gal80. Null MARCM clones were generated using the following stocks: Odd^rk111^line FRT40A (kind gift of Cordelia Rauskolb) (Hao, Green et al. 2003), hsFLP, UAS-mCD8::GFP. tubP-GAL80 neoFRT40A; tubP-GAL4. We used the UAS-odd-skipped for all miss and over expression study. The following FlyLight (Janelia Farm) Gal4 stocks were used for miss expression GMR69G07 and GMR17B12 as well as Elav-Gal4. Bruchpilot tagged with GFP is a MiMIC insertion (Nagarkar-Jaiswal, Lee et al. 2015).

### Immunofluorescence microscopy

Adult and larval brains were dissected in cold phosphate-buffered saline (PBS; pH 7.2) and fixed in PBS-buffered 4% formaldehyde for 30 minutes. Brains were washed several times in PBS and stored in methanol at 20 °C. Standard antibody labelling was followed (Ashburner, 1989). Briefly, brains were rehydrated in PBS containing 0.5% Triton X-100 (Sigma; PBS/Triton) and incubated in PBS/Triton plus 10% goat serum (Sigma) for 3 hours. Brains were incubated with primary antibody for 2 days and secondary antibody overnight. The brains were washed several times in PBS/Triton between the primary and secondary antibody. For immunohistochemistry on embryos, flies were allowed to lay eggs on yeasted apple juice agar plates for 12 hours. Embryos were collected, dechorionated in bleach, and fixed in PBS-buffered 4% formaldehyde. Antibody staining was performed as for adult and larval brains except Tween-20 (Sigma) was used instead of Triton X-100. Also, primary antibodies were incubated overnight and secondary antibodies for 2 hours.

Immunohistochemistry using Odd-skipped antibody differed from the standard protocol as follows: brains were fixed using PLP fixative (Brenes, Harris et al. 1986) and incubated in PBS/Triton plus 10% goat serum and 0.5% deoxycholate overnight at 4 °C. Primary antibody solution was also supplemented by 0.5% deoxycholate. Brains were mounted in PBS and viewed using a Zeiss 510 confocal microscope using either an air 20x or oil-immersion 40x objectives.

### Generation of Odd-skipped antibody

Odd-skipped antibody is a polyclonal guinea pig antibody, produced by DundeeCell products (www.dundeecellproducts.com). The following peptides were used for antibody production: CSTSASPISNITVD and CGASGVPSGATGS which localises to start and middle of the Odd-skipped polypeptide (in green); MSSTSASPIS NITVDDELNL SREQDFAEED FIVIKEERET SLSPMLTPPH TPTEEPLRRV HPAISEEAVA TQLHMRHMAH YQQQQQQQQQ QQQHRLWLQM QQQQQQHQAP QQYPVYPTAS ADPVAVHQQL MNHWIRNAAI YQQQQQQQQH PHHHHHHGHP HHPHPHPHHV RPYPAGLHSL HAAVMGRHFG AMPTLKLGGA GGASGVPSGA TGSSRPKKQF ICKYCNRQFT KSYNLLIHER THTDERPYSC DICGKAFRRQ DHLRDHRYIH SKDKPFKCSD CGKGFCQSRT LAVHKVTHLE EGPHKCPICQ RSFNQRANLK SHLQSHSEQS TKEVVVTTSP ATSHSVPNQA LSSPQPENLA QHLPVLDLSS SSSSSEKPKR MLGFTIDEIM SR. The antibody was isolated from antisera by immunoaffinity chromatography using antigen coupled to agarose beads. For wholemount immunohistochemistry the Odd-skipped antibody was used at a 1:500 dilution.

### Antibodies

The following antibodies were used: mouse monoclonal anti-Bruchpilot (nc82;1:10 dilution; Developmental Studies Hybridoma Bank), polyclonal rabbit anti-GFP (1:500 dilution; Invitrogen) and polyclonal chicken anti-galactosidase (1:1000 dilution; catalog #ab9361, AbCam). The Syndapin antibody (1:100 dilution, kind gift from Vimlesh Kumar) (Kumar, Fricke et al. 2009). The secondary antibodies (Invitrogen) used were as follows: Alexa Fluor 488 donkey anti-rabbit, Alexa Fluor 546 donkey anti-chick, and Alexa Fluor 546 goat-anti-mouse. These were used at a 1:500 dilution. For live imaging of Bruchpilot tagged with GFP larval brains were dissected in cold PBS, mounted, and viewed as for fixed preps.

### Western blots

Brains were dissected in PBS and immediately frozen in an Eppendorf tube on Dry Ice. After defrosting samples were immediately homogenised in sample buffer containing 80mM Tris pH6.8, 0.1M DTT, 10% glycerol and 2% SDS. Samples were denatured at 95°C for 10min and protein quantity was measured using Pierce 660nm Protein Assay (ThermoFisher). Protein extracts (10µg) were separated on 10% acrylamide gels for 2h at 120V and transferred onto methanol-activated PVDF membranes at 350mA for 2h. Membranes were blocked with 5% non-fat milk (Biorad) in TBS-T (20mM Tris-HCl pH7.5, 150mM NaCl and 0.1% Tween20) for 1h and probed with the following primary antibodies: guinea-pig anti-Oddskipped (1/100, in house) and mouse anti-actin-HRP (1/20 000, Sigma A3854). After incubation with HRP-conjugated anti-guinea-pig secondary antibody (1/10 000, ThermoFisher PA1-28679), protein levels were visualised by chemiluminescence with an Odyssey FC (Li-Cor) and quantified with Image Study Lite. For quantification, densitometry of the band of interest was normalised to that of actin.

### Chemotaxis assay

The chemotaxis assay has been described previously (Slater, Levy et al. 2015). Briefly larvae were collected, washed, and transferred onto a 1% agarose Petri dish followed by 2-hour starvation. All assays were performed at room temperature and to avoid external cues, such as light, affecting larval behavior, we always performed the assays at the same location and always rotated the behaviour arena 90° between each experiment. Approximately 50 early second-instar larvae were placed in zone 0 (see Fig. 9A) on a 1% agarose gel. 20 µl of Apple cyder vinegar was added to a 0.5 cm Eppendorf lid placed in zone +4 directly opposite each another lid containing 20 µl of H_2_O placed in zone ࢤ4. Immediately after adding odor to the arena, a lid containing 7 ventilation holes was placed on top of the Petri dish, and the larvae were allowed to wander for 5 min. At the end of the experiment, the number of larvae in each zone was calculated. Larvae that did not leave zone 0 were not included in the data collections. A response index (RI) was calculated by deducting the number of larvae in a given minus zone from that of the number of larvae in the equivalent plus zone divided by the total number of larvae in the assay. A positive RI shows that larvae can chemotax toward the odor (zone +4), whereas an RI of 0 would indicate that larvae have no preference for the odor over the control side. A negative RI shows that larvae are repelled by specific odors.

### Live imaging and locomotion tracks

We imaged larval behaviour during the chemotaxis assay. Larvae were imaged using a Nikon SMZ1500 dissecting microscope with ambient light at room temperature. Movies were captured on a Nikon Digital sight DSRi1 camera at a frame rate of 7 frames/s. FIMtrack software (Risse, Thomas et al. 2013) was used to analyse turning behavior during the chemotaxis assay. As part of the analysis, FIMtrack generates full locomotion tracks of all the larvae that were analyzed. We combined the locomotion tracks from all the analysed larvae in CorelDRAWX3(Corel) and manually traced over the tracks.

### Statistical and behavioural analyses

GraphPad Prism5 was used for all statistical analyses. Each chemotaxis assay was repeated 10 times, and statistical significance was calculated using a two-way ANOVA test and Bonferroni post-test. FIMTrack was used to analyse the frequency of larval body bending to quantify number of head swings/minute. We pre-set the angle to 60° in the software so that body bending at a lower degree was not included in the analysis. A Kruskal–Wallis test combined with a Bonferroni post-test was used to analyse statistical significance among the genotypes tested. Fluorescence tagged antibodies were used to visualise both Bruchpilot and Syndapin protein levels in Odd neurons and neighbouring tissue (e.g. Fig. 8G). Fiji64 (ImageJ) was used to quantify fluorescence intensity using ROI manager. Statistical significance between Odd neurons and neighbouring tissue was analysed using a 2-way ANOVA with a Bonferroni post-test. Fiji64 ROI manager was also used to quantify GFP fluorescence intensity between Odd neural arbours and adjacent tissue of Bruchpilot tagged with GFP. Statistical significance was calculated using a 2-way ANOVA and a Bonferroni test. A one-way ANOVA test was used to calculated statistical significance between genotypes for the Western blots.

## Results

### Loss-of-Odd function causes stunted neurite growth

To begin to understand the role of Odd in central brain neurogenesis we first took a loss of function approach. Loss-of-function of Odd is embryonic lethal (Coulter and Wieschaus 1988) and we therefore adopted the MARCM approach (Lee and Luo 2001) to induce Odd null clones at later developmental stages. The MARCM approach also allowed us to describe morphological changes in more detail since axonal and dendritic morphology is more complex at larval and adult stage compared to the embryo. Clones induced at larval stage (48 and 96 hours after egg lay) (Fig 1A-F) were analysed at adult stages whereas clones induced during embryogenesis (8 hours after egg lay) (Fig 1G,H) were analysed at 2^nd^ instar larval stage. To allow us to differentiate between clones generated in Odd expressing cells and clones induced in non-Odd expressing neurons we took advantage of a null allele of *Odd-skipped* with an insertion of LacZ in the coding region of the Odd gene (Hao, Green et al. 2003). The expression of LacZ in this line replicates the normal expression of Odd in the brain (Levy and Larsen 2013). Clones induced in the Odd expressing neurons will therefore co-express both GFP (expressed in neurons that have two copies of the null *Odd-skipped* allele) and LacZ. Regardless of developmental stage of clonal induction all *Odd-skipped* null clones exhibit the same morphology (Fig 1A-H).

**Figure 1.**
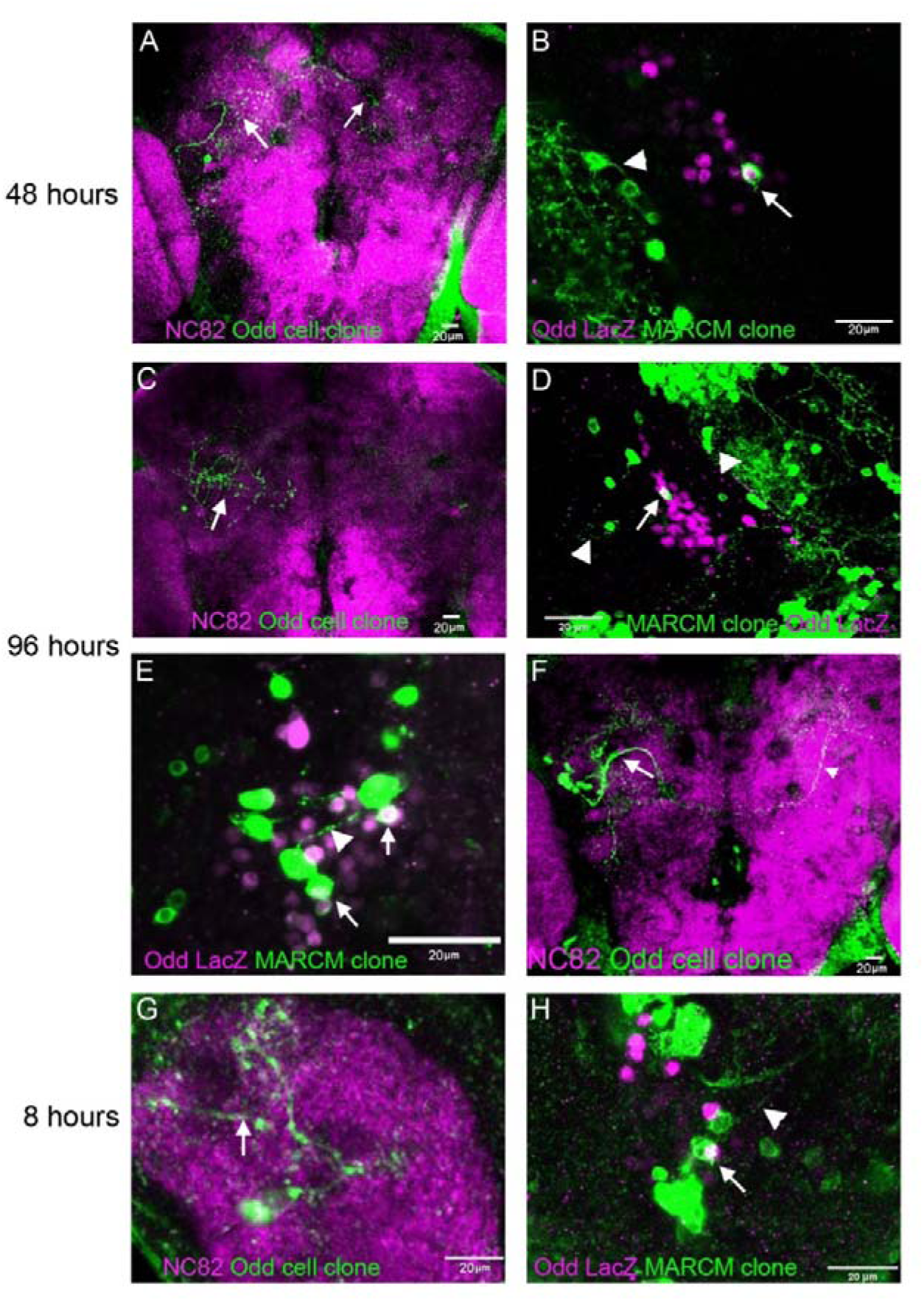
Loss-of-Odd function inhibits neurite outgrowth. Dorsal view of the brain, anterior to the top, posterior at the bottom. A-B) Clones induced at 48 hours after egg lay. A) Wildtype clone demonstrating that clonal induction at 48hours labels a neuron which projects into the several compartments and projects across the midline arborizing in the contralateral side of the brain (arrows). B) A MARCM null clone induced at the same developmental stage. The neurite is stunted (arrow), whereas clones induced in non-Odd expressing neurons all have long neurites (arrowhead). C-F). Clonal induction at 96 hours after egg lay. C) A wildtype clone showing elaborate neurite morphology (arrow). D) There is no neurite visible in the MARCM null clone (arrow) whereas neighbouring non-Odd expressing neurons exhibit elaborate neurite morphology (arrowhead). E) Another example of a MARCM null clones lacking neurites (arrows) whereas a neighbouring non-Odd expressing clone extends a neurite medially (arrowhead). F) Induction of wildtype clone can also induce a lineage clone demonstrating that induction the MARCM approach used in this study can induce NB clones which extend neurites both ipsi and contralaterally (arrows). G-H) Induction of MARCM and wildtype clone at 8 hours after egg lay. G) Induction of wildtype clone at 8 hors labels a cell the project into the calyx of the MB and innervate the CPM compartment (arrow). H) The MARCM clone induced at a similar developmental stage has a small protrusion (arrow) whereas neighbouring non-Odd expressing cells have long neurite extensions.

Neurites are either missing all together (Fig 1D,E) or appear as small stunted protrusions (Fig 1B,H). Wildtype clones induced at the same developmental stage (Fig 1A,C,F,G) all exhibit a large arbour morphology showing neurite projections into multiple areas of the brain (Fig 1A,C) and projections cross the midline which arborize in the contralateral side of the brain (Fig 1F). The change in neurite morphology of Odd null clones is unlikely to be due to an unknown genetic background effect since non-Odd expressing neurons in the same brain exhibit longer neurites with elaborate morphology (Fig 1B, D, H). This data show that loss of Odd function results in a deficit in neurite formation suggesting that Odd could play a role in outgrowth and extension. In addition, we never observed multi-cell Odd expressing clones whereas control clones induced at a similar developmental time often contained multiple labelled cells. This suggests that lack of Odd in the neuroblast (NB) either terminates NB division or affects survival of the NB.

### RNA interference causes neural overgrowth

Due to the severity of the Odd null phenotype we wondered whether reducing Odd protein levels would lessen the severity of the phenotype. This would allow us to gain a better understanding of which aspect of neurite growth is controlled by Odd. To achieve this, we used an RNAi approach initially investigating the effect of expressing 3 different RNAi lines; one generated by the Vienna Drosophila resource centre (VDRC) (ref) and 2 from the TRiP project, expressed in Odd expressing cells using the Odd-Gal4 driver line. We included UAS-CD8:GFP in all our genetic crosses allowing us the characterise the neurite morphology of Odd neurons. Heterozygous expression of RNAi against Odd results in a similar phenotype regardless of RNAi construct used, although the VDRC RNAi line phenotype is more severe (Fig 2A-L). RNAi expressing Odd neurons are bushier and denser and, in some cases, extend into brain regions that are not innervated by wildtype Odd neurons (Fig 2 A-D). For example, RNAi expression affect the neurites that project towards the AMMC which are thicker and have a bushier appearance (Fig 2D) compared to wildtype Odd neurons (Fig 2C, K). The projections from wildtype Odd neurons are tightly fasciculated and it is possible that the phenotype seen in the Odd neurons expressing the RNAi is caused by de-fasciculation. A similar “de-fasciculation “phenotype is seen in the neurites that cross the midline (Fig 2D,J,L) which are thicker and bushier compared to the tightly fasciculated neural track of the wildtype Odd neurons (2C,K). Neurites also appear to extend into regions of the brain in which wild type Odd neurons do not project (Fig 2A-B). This is particularly evident in Odd neurons expressing the VDRC RNAi and to a lesser extend for the trip 34328 RNAi line. We have previously described the projection pattern of Odd neurons in the brain (Levy and Larsen 2013) and here we adopt the terminology of the Insect brain name working group. Wildtype Odd neurons innervate medial part of the PVLP (Fig 2C, K) whereas Odd neurons expressing both the VDRC and trip 34328 RNAi innervate most of PVLP (Fig 2D, L). VDRC expressing Odd neurons extend into the SCL, SIP and SMP regions (Fig 2B) which are not innervated by wildtype Odd neurons (Fig 2A). Wildtype Odd neurons extend a single neurite into the SPS (Fig 2C). Expressing the VDRC RNAi in Odd neurons results in extensive projections into this region (Fig 2D) with projections extending into the GNG. Expressing the trip 28295 RNAi produced the weakest phenotype (Fig 2E-H) and only seems to affect the projections towards the AMMC (compare Fig 2E and F) and the neurites crossing the midline (compare Fig 2G and H). Thus, it would appear that reducing Odd protein expression levels have the opposite phenotype to that off the null phenotype. Only the lobular plate projections appear to show a reduction in arbour growth (Fig 2N) compared to wildtype Odd neurons (Fig 2M) and 1 projection fail to extend into the lobular plate altogether terminating in the lobular (Fig 2N).

**Figure 2.**
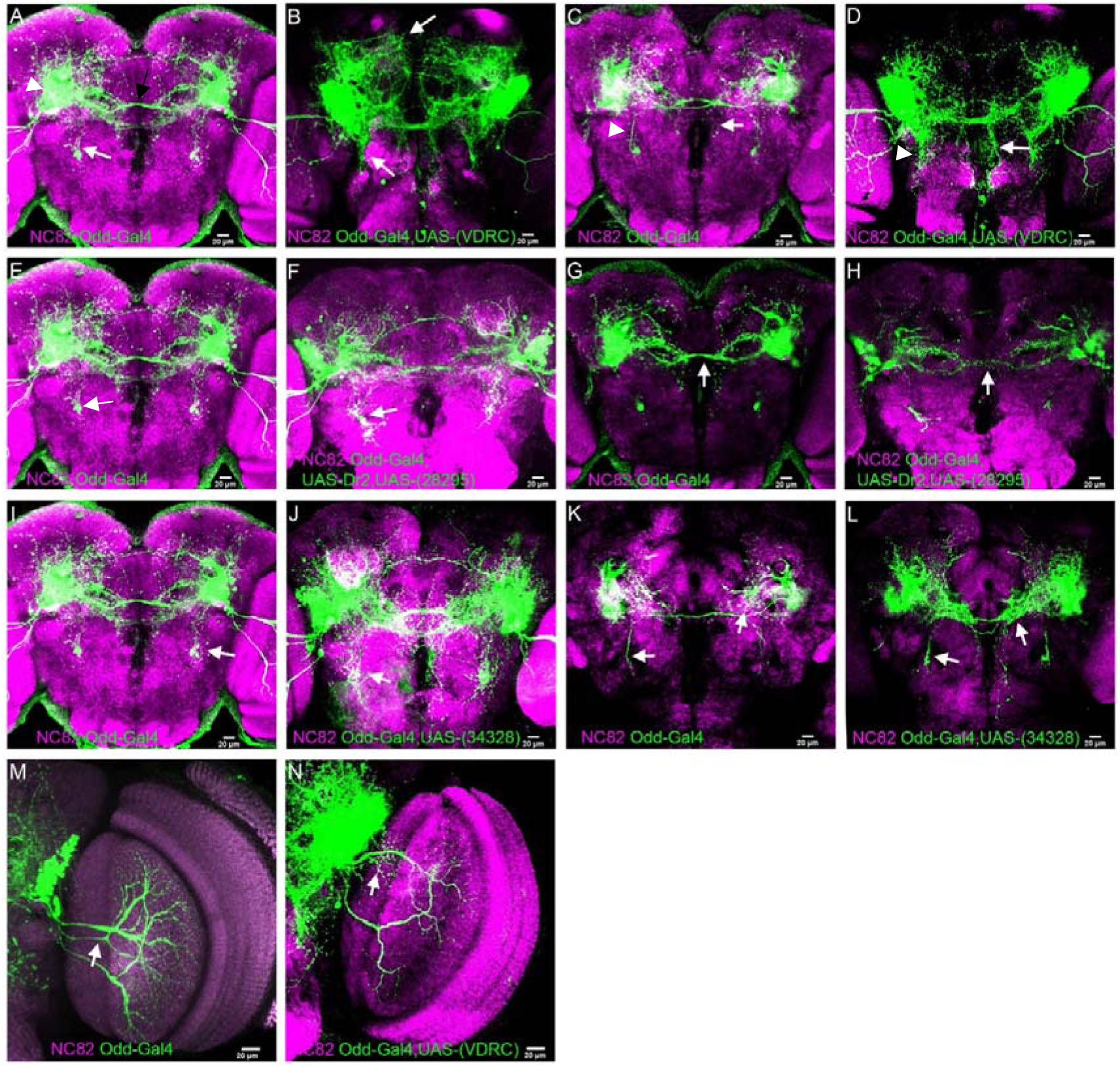
Expressing RNAi against Odd causes neural overgrowth in the adult. Dorsal view of the brain, anterior at the top, posterior at the bottom. A-D. Expressing of the VDRC RNAi against Odd. A,B,E,F,I,J are full z projections and C,D,G,H,K,L are single sections at a similar dorso-ventral position in the brain. A) Wildtype Odd neurons project into part of the PLP (arrowhead) and towards the AMMC (white arrow). They also extend tightly fasciculated projections across the midline (black arrow). B) Odd neurons expressing the VDRC RNAi extend projections SCL, SIP and SMP regions of the brain (anterior arrow) and the projection towards the AMMC is thicker and bushier (posterior arrow). C) A single section of the z-stack shows the single wildtype Odd neural projection into the SPS (arrow) as well as the tightly fasciculated AMMC projection (arrowhead). D) A single section of the z-stack shows the increase in projections into the SPS (arrow) and the bushier appearance of the AMMC projection (arrowhead) of the Odd neurons expressing the VDRC RNAi. E-H) Comparison of Odd neural projection between wildtype and Odd neurons expressing the TRiP RNAi line 28295. E) Z-stack of wildtype Odd neural projections highlighting the projection towards the AMMC (arrow). F) Z-stack of RNAi 28295 expressing Odd neurons. The phenotype is less severe than that of VDRC expression but the AMMC projections also appear de-fasciculated when expressing RNAi 28295 (arrow). G) Single section of z-stack of wildtype Odd neurons showing the morphology of the midline crossing neurites (arrow). H) A single section of the z-stack shows that the midline crossing neurites are less fasciculated in brains expressing RNAi 28295 (arrow). I-L) Comparison of Odd neural projection between wildtype and Odd neurons expressing the TRiP RNAi line 34328. I) Z-stack of wildtype Odd neural projections highlighting the AMMC projection (arrow). J) Z-stack illustrating the highly de-fasciculated AMMC projections in brains expressing RNAi 34328 (arrow). K) Single section of the z-stack showing the Odd neurons do not project into the posterior part of the PLP (arrow) whereas L) Odd neurons expressing the RNAi 34328 innervate the entire posterior part of the PLP (arrow). M) 3 of the Odd neurons project into the lobular plate of the optic lobe (arrow). N) One of the Odd neurons fail to innervate the lobular plate in brains expressing RNAi 34328 in the Odd neurons. Instead this neuron appears to innervate the lobular immediately adjacent to the central brain (arrow)

### Homozygous expression of RNAi disrupts neurite arbour formation and midline crossing

We were puzzled by the apparent opposite phenotype generated by the expression of RNAi in Odd neurons. We hypothesised that this could be due to insufficient lowering of Odd protein expression by only expressing 1 copy of the RNAi. We therefore generated double homozygous flies with 2 copies of Odd-Gal4 and UAS-RNAi thereby theoretically doubling the level of RNAi produced in the cell. Homozygous Odd-Gal4, UAS-RNAi is larval lethal for all RNAi lines tested and larvae die at the end of 2^nd^ instar. Since the Odd-Gal4 used in this study drives expression in all Odd expressing cells it is highly likely that the lethality is not due to the expression in Odd neurons but more likely due to the expression in other tissues that also express Odd such as the epidermis. Given the larval lethality we therefore compared Odd neural projection patterns of RNAi homozygous brains to that of similar staged heterozygous and wildtype brains (Fig 3A-I). In the wildtype larvae Odd neurons project dendrites into the calyx and extend axonal projections both ipsi- and contralateral arborizing in the CPM (Fig 3A). Since the RNAi expressing larvae are raised at 29 C° we also raised control larvae at this temperature. Due to the transient expression of Odd in 2 of the Mushroom Body NBs during embryogenesis the temperature induced increase in GFP expression and perdurance of GFP, some GFP positive MB neurons are still visible at 2^nd^ instar larval stage (marked by * in Fig 3A). At 2^nd^ instar larval stage heterozygous expression of all 3 RNAi lines results in little disruption of arbour formation (Fig 3B,D,F) compared to wildtype arbour morphology (Fig 3A) with the exception that the midline crossing neurites appear de-fasciculated. Homozygous expression on the other hand, disrupts midline crossing (Fig 3C, E, G) and the axonal arbours localised in the CPM also appear disorganised and are no longer localised to the CPM exclusively. Dendritic projections into the calyx are reduced or absent (Fig 3C, E, G). Because all three RNAi lines produce similar phenotypes, we decided to progress with the VDRC line and the trip 34328 line as the latter line produced consistently stronger phenotypes compared to the trip 28295 line.

**Figure 3.**
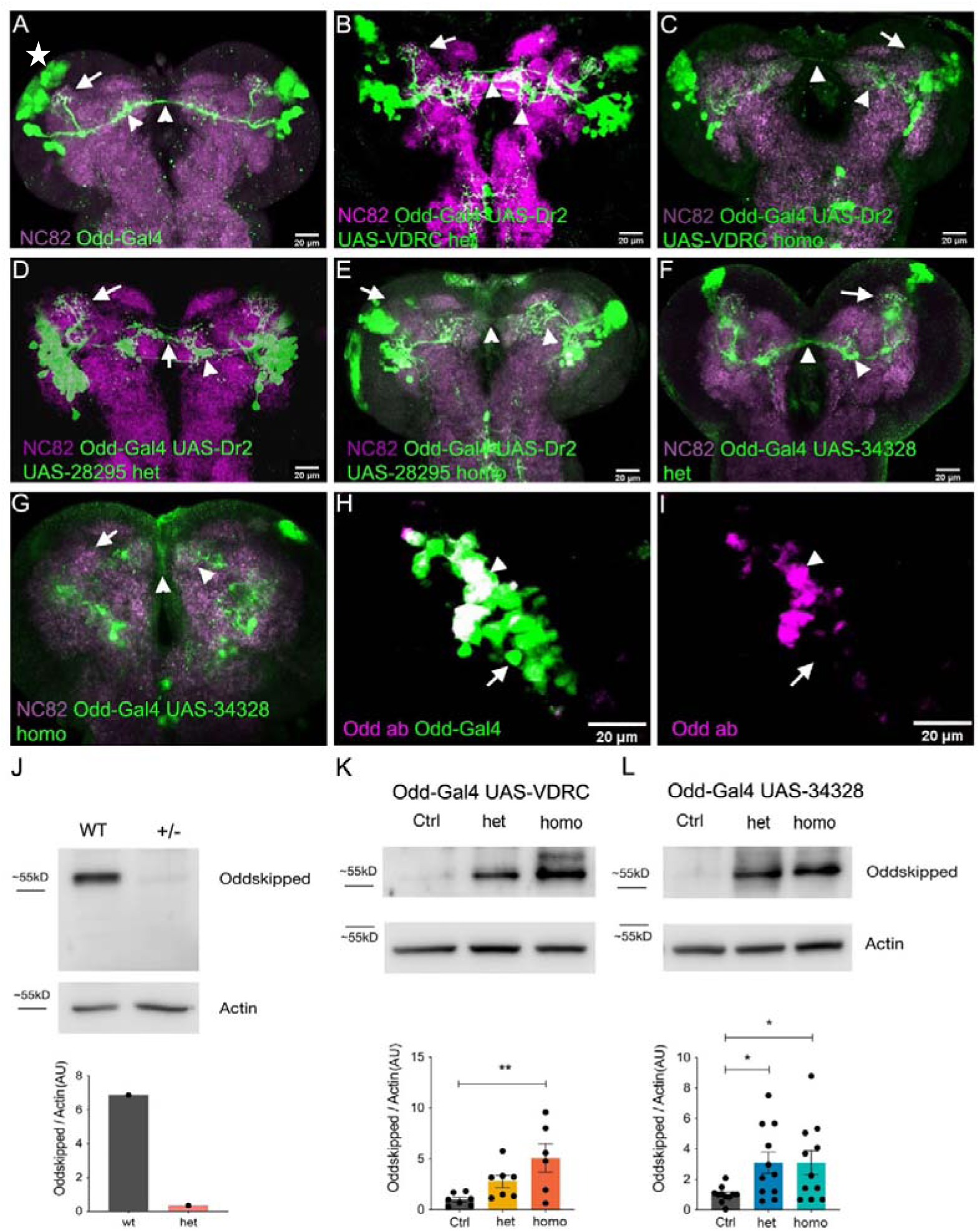
Homozygous RNAi expression disrupts neurite growth and arborisation. Dorsal view of the brain, anterior at the top, posterior at the bottom. A) Wildtype Odd neurons project dendrites into the calyx (arrow) and axons across the midline which arborize in the contralateral CPM compartment (arrowheads). B-C). Hetero- and homozygous expression of RNAi-VDRC in Odd neurons respectively. Heterozygous expression has little effect on Odd neurite growth and arborisation. The dendritic arbour in the calyx is normal (arrow) but the midline crossing does appear defasciculated (arrowhead). Homozygous expression results in a larger axonal arbour and a reduction in axonal midline crossing (arrowhead). Dendritic projections into the calyx are gone (arrow). D-E). Hetero- and homozygous expression of RNAi 28295 in Odd neurons respectively. Heterozygous expression does not affect axonal and dendritic projections (arrow and arrowhead) whereas homozygous expression abolishes projections into the calyx (arrow) and axonal midline crossing (arrowhead). The axonal arbour is spread out (arrowhead). F-G) Hetero- and homozygous expression of RNAi 34328 in Odd neurons respectively. Heterozygous expression has no affect on Odd neural arborisation in the calyx (arrow) and axonal projections (arrowhead), whereas homozygous expression severely disrupts axonal arbour formation (arrowheads). There is no axonal midline crossing and the axonal arbour is reduced and spread out. The dendritic projections into the calyx are absent (arrow). H-I) Odd Antibody (Kumar, Fricke et al.) co-localisation with GFP expressed in the Odd neurons using the Odd-Gal4 line. Many of the GFP expressing neurons co-localises with Odd antibody (arrowhead) but some GFP positive neurons do not (arrow). J) Western blot showing Odd protein levels in wildtype brains compared to brains heterozygous for the Odd null allele (Odd^rk111^). Odd protein levels are reduced in heterozygous Odd^rk111^ brains. K) Western blot of Odd protein levels in control brains (Odd-Gal4), heterozygous and homozygous expression of RNAi VDRC. Odd protein levels are significantly increased (1 way-ANOVA) in homozygous RNAi VDRC expressing brains compared to control and heterozygous expressing brains. L) Western blot of Odd protein levels in control brains (Odd-Gal4), heterozygous and homozygous expression of RNAi 34328. Odd protein levels are significantly increased (1 way-ANOVA) in homozygous and heterozygous RNAi 34328 expressing brains compared to control.

H0mozygous expression of RNAi constructs have a stronger phenotype that heterozygous expression which could be due to differences in Odd protein levels presumably being lowered by the additional copy of Odd-Gal4 and UAS RNAi. To confirm that the difference in phenotype was due to different levels of Odd protein we generated an antibody against Odd in order to quantify protein levels using western blots. First, we tested the specificity of the Odd antibody using wholemount immunohistochemistry by analysing co-localise between Odd antibody and the GFP expression driven by the Odd-Gal4 driver. It should be noted that in wholemount the Odd antibody does not produce a strong stain. The majority of GFP labelled Odd neurons co-labels with Odd antibody (Fig 3H, I) although some GFP positive cells are not labelled by the Odd antibody. This could be due to low levels of Odd expression in these neurons combined with less optimal binding of the antibody. Importantly we did not observe Odd antibody labelling of other neurons in the brain suggesting that the Odd antibody is specific to Odd. We next sought to verify that the antibody was specific for Odd in a western blot by comparing Odd protein levels between wildtype larval brain and the heterozygous null allele Odd-LacZ. The Odd antibody binds to a 55KD protein (Fig 3J) and is strongly reduced in the heterozygous Odd-LacZ brains confirming that the antibody likely is specific to Odd. We next analysed Odd protein levels in the larval brains either heterozygous or homozygous for the VDRC RNAi and found to our surprise that RNAi expression significantly upregulates Odd protein levels (Fig 3K). Homozygous expression increases Odd expression levels more strongly that heterozygous expression. A similar result is obtained with the TRIP 34328 line although both heterozygous and homozygous expression upregulates Odd protein to a similar level (Fig 3L). This data shows that upregulating Odd protein levels affect neurite growth and arborisation and clearly demonstrates that RNAi can increase proteins levels. Unfortunately, in this case, RNAi against Odd cannot be used to lower Odd protein levels.

### Overexpression of *Odd-skipped* results in a similar phenotype to that of RNAi expression

Our RNAi data suggests that increasing Odd protein levels affect neurite growth and arborisation. We would therefore predict that over-expressing Odd should result in a similar phenotype to that of RNAi expression. In order to address this, we expressed Odd in Odd neurons using the Odd-Gal4 driver. Heterozygous expression of Odd is viable but homozygous expression is embryonic lethal using 2 copies of Odd-Gal4 and UAS-Odd. However, we found that flies expressing 1 copy of Odd-Gal4 and 2 copies of UAS-Odd survived into larval stages and like the homozygous RNAi expressing flies died at the end of 2^nd^ instar larval life. Heterozygous expression of Odd in the Odd neurons has a similar affect to that of driving heterozygous expression of Odd RNAi although the phenotype is not as strong (Fig 4A-D). For example, the neurites projecting towards the AMMC are bushy and the neurites crossing the midline are disorganised (Fig 4A-D). Otherwise there is little effect on projection patterns. Prior to analysing the effect of expressing 2 copies of UAS-Odd we first wanted to gain an understanding of how over-expression of Odd affects Odd protein levels using western blots. Odd Over-expression does course upregulation of Odd protein levels compared to the control demonstrating that Odd protein is upregulated using the over expression approach (Fig 4E). In the larvae heterozygous expression of Odd does not affect midline crossing or dendritic arborisation in the calyx (Fig 4G) but the CPM arbour appears to be larger than in control larvae (Fig 4F). Odd-Gal4 with 2 copies of UAS-Odd affects fasciculation of the neurites crossing the midline (Fig 4H) and CPM arbours are reduced as is the dendritic innervation of the calyx although some dendritic branching is present. Thus, the phenotype matches that of RNAi although appears not to be as severe. Due to the obvious de-fasciculation phenotype observed in both over-expression and RNAi expressing Odd neurons we sought to address the effect of increasing Odd protein levels on the expression of genes known to be involved in fasciculation in *Drosophila*, including NCAD, Connetin, FasII and Semaphorin 1a. The expression levels of these genes are not affected by Odd over expression (data not shown) either using RNAi or UAS-Odd.

**Figure 4.**
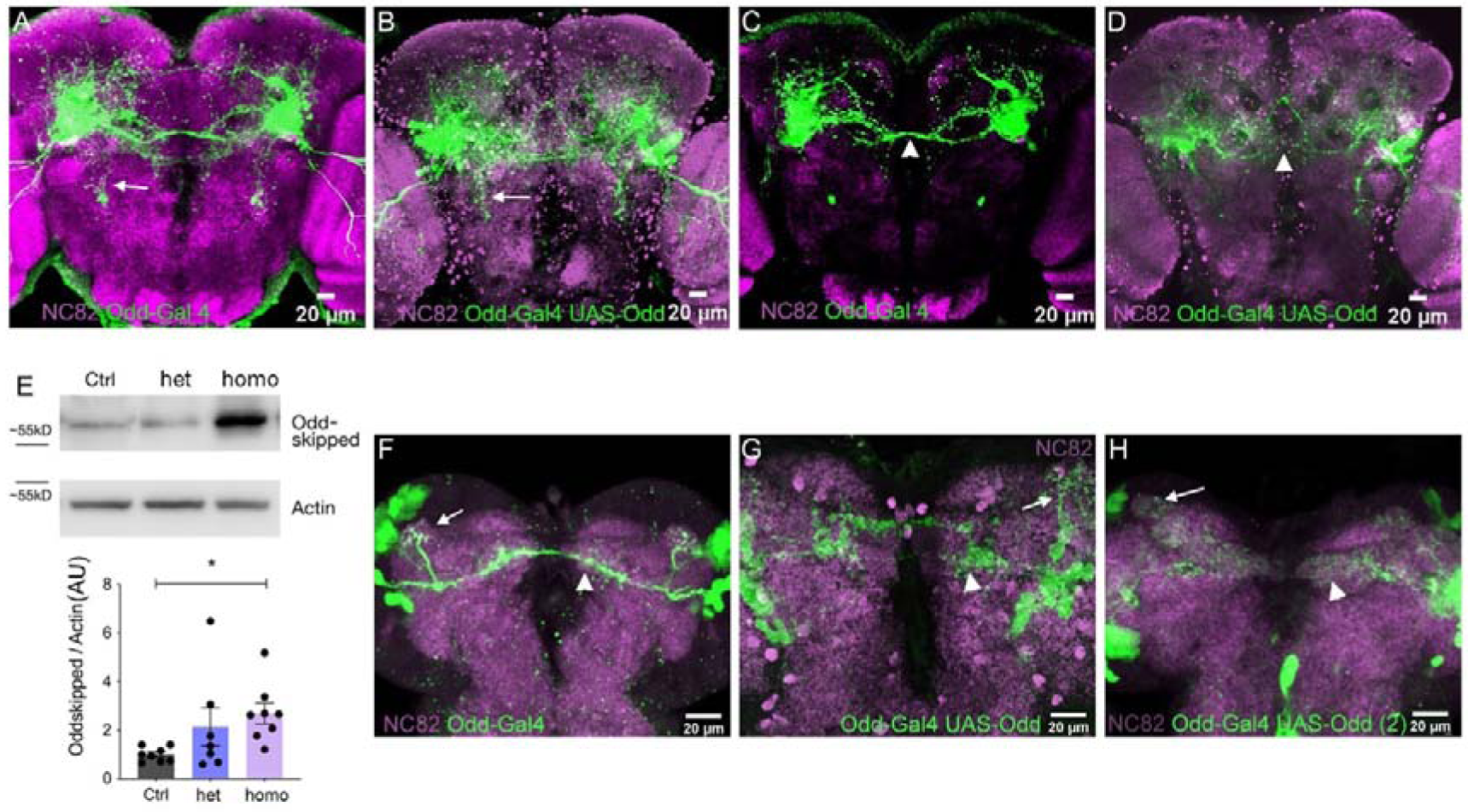
Expressing Odd in the Odd neurons result in a similar phenotype to that of RNAi expression. Dorsal view of the brain, anterior at the top, posterior at the bottom. A) Z-stack projection showing wildtype Odd neural projection highlighting the AMMC projections (arrow). B) Z-stack projection of Odd neurons expressing UAS-Odd. The AMMC projection is defasciculated (arrow). C-D) Single section of the z-stack shown in A and B respectively at the same dorso-ventral level of the brain. Wildtype Odd neurons form tightly fasciculated tracks across the midline (arrowhead), whereas midline crossing appears defasciculated in the brains which express UAS-Odd in the Odd neurons. E) Western blot comparing Odd protein levels between control (Odd-Gal4), hetero (Odd-Gal4 UAS-Odd) and homo (Odd-Gal4 UAS Odd 2). Expressing 2 copies of UAS-Odd significantly increased (1 way-ANOVA) compared to controls and expression of 1 copy of UAS-Odd. F) Wildtype Odd neurons project dendrites into the calyx (arrow) and axons across the midline which arborize in the contralateral CPM compartment (arrowheads). G) Expressing 1 copy of UAS-Odd does not affect dendritic projections (arrow) but the axonal arbour does appear defasciculated (arrowhead). H) Expressing 2 copies of UAS-Odd affects dendritic projections to the calyx, although some branches are visible within the calyx (arrow), but axonal projections are defasciculated (arrowhead).

### Miss-expression of *Odd-skipped* causes defects in neurite growth and arborisation and neural lethality

Next, we asked whether Odd affects neurite growth and arborisation in neurons that do not normally express Odd in order to understand if Odd plays a fundamental role in neurite formation. We started addressing this question by expression Odd in all post mitotic neurons using the ELAV-Gal4 driver. ELAV is exclusively expressed in the CNS in *Drosophila (Yao and White 1994)* and therefore allows us to target expression of Odd specifically to neurons. We co-express CD8-GFP to visualise all neurons and their projections. It is therefore surprising that expression of Odd using the ELAV-Gal4 driver is embryonic lethal. In the embryo the size of the brain appears normal compared to control (Fig 5A-B). However, the number of midline crossing fascicles are reduced in brains in which Odd is expressed in all neurons compared to the control (ELAV-Gal4) (Fig 5C-G). This is particularly evident when examining single optical sections (Fig 5E, F) of a z-stack at the same position in the brain. Control embryos have 3 distinctive tracks crossing the midline whereas embryos expressing Odd only has 1 track crossing the midline. Across multiple brains all 4 tracks were always present in control brains whereas the number of tracks varied from 1 to 3 in brains expressing Odd (Fig 5G). This data would suggest that mis-expressing Odd affects neurite growth but only in a subset of neurons since some neural tracks remain in the brain in which Odd is miss-expressed. Furthermore, similar to Odd over expression using either RNAi or UAS-Odd the severity of the phenotype varies.

**Figure 5.**
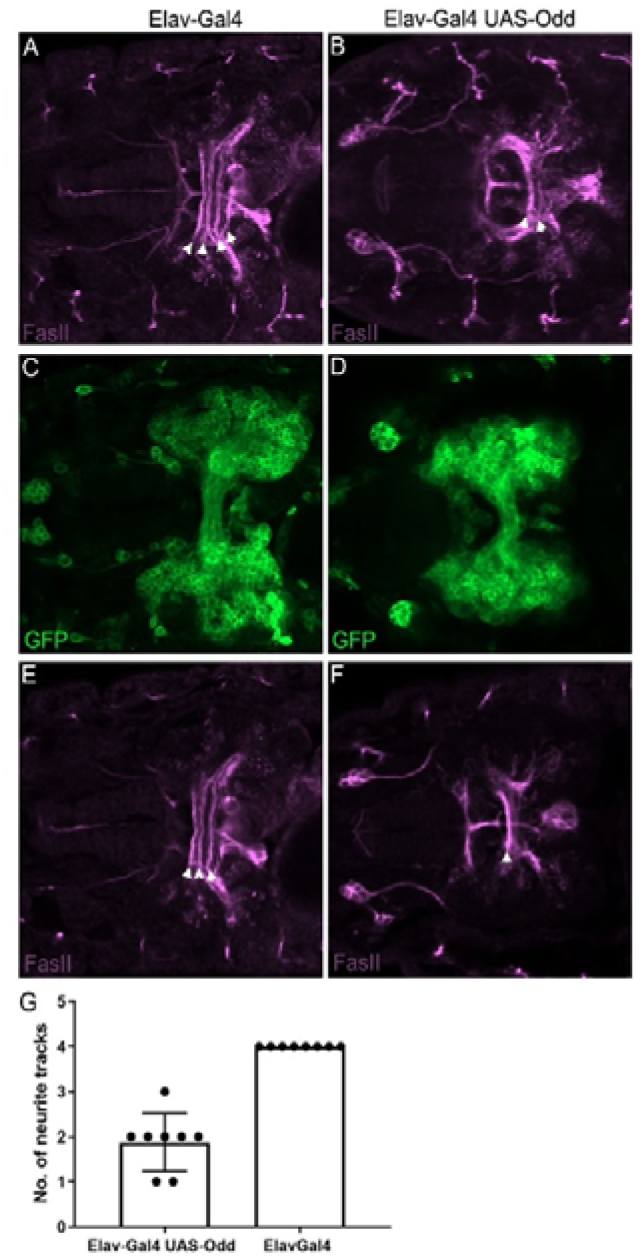
Miss-expressing Odd in all postmitotic neurons reduces nerite midline crossing. Dorsal view of the brain. Anterior to the right. Scale bars=10 µm A-B) Z-stack projection of the entire brain of control (Elav-Gal4) and expression of Odd using the Elav-Gal4 driver respectively. FasII labels neurite tracks. 4 neural tracks are present in the control brain (arrowheads) whereas brains expressing Odd only have 2 midline crossing tracks (arrowheads). C-D) Z-stack projection of the entire brain of control (Elav-Gal4) and expression of Odd using the Elav-Gal4 driver respectively. Brain morphology does not appear to be affected miss-expression of Odd. E-F) Single section of the Z-stack projection shown in A and B at a similar dorso-ventral position in the brain. At this level of the brain 3 tracks are present in control brains whereas brains in which Odd is miss-expressed only have 1 track. G) Number of track present in control brain and Odd miss-expressing brains.

To explore this further we used a number of different Gal4 driver lines from the FlyLight collection (Tokusumi, Tokusumi et al. 2017) to selectively express Odd in small subsets of neurons (Fig 6A-H). The phenotype of miss-expressing Odd across different driver lines supports the hypothesis that both Odd protein levels and neural subtype determines the severity and type of the phenotype produced. For example, using the R69G07-Gal4 line shows that miss-expressing Odd results in different phenotypes depending on neural subtype. The R69G07 line is expressed in two sets of neurons (Fig 6A-B). One neuron in each hemisphere send projections anteriorly and arborize in several compartments laterally in the brain whereas a separate small group of neurons arborize exclusively in the medial lobe of the MB. The neurons that project to the MB are missing in brains in which Odd is miss expressed (Fig 6B) whereas the arbours arising from the single neurons located more anteriorly are maintained although neurites do not extend into the lateral part of the brain. A similar phenotype is seen using the driver line R17B12-Gal4 (Fig 6C-J). This Gal4 line drives expression in a cluster of neurons situated anteriorly in the brain which project posteriorly and cross the midline (Fig 6C). These neurons are missing in brains in which Odd is miss-expressed although in approximately 1:11 brains, the neurons survive in the left hemisphere (Fig 6D). In those cases, neurite growth is severely stunted and appear to emerge randomly from the cell. The R17b12-Gal4 also drives expression in the laminar neurons in the optic lobe (Fig 6E). In control brains laminar neurons project into the medullar (Fig 6E) whereas laminar neurons miss-expressing Odd do not exhibit in obvious neurite growth (Fig 6F). The laminar cells bodies also appear to be disorganised compared to the control neurons which are tightly packed (Fig 6E). Our data show that miss expressing Odd skipped affects neurite growth and arborisation. In addition, miss expression may also affect cell survival or induce cell death. To explore this further we first wanted to establish whether the neurons that are missing in larval brains are present at embryonic stage and if so whether neurite outgrowth/arborisation is affected. For all the Gal4 lines in which neurons are missing at larval stage we saw driver line expression in the embryo, showing that the neurons are present at early developmental stages. For example, R17B12-Gal4 drives expression in neurons localised to the posterior part of the embryonic brain which extend projections anteriorly and across the midline (Fig 6G,H). Expressing Odd in these neurons cause a failure in neurite midline crossing and neurite projections terminate close to the cell bodies (Fig 6I,J). Thus, miss expression of Odd does not affect neurogenesis but specifically affects neurite outgrowth/arborisation prior to cell death.

**Figure 6.**
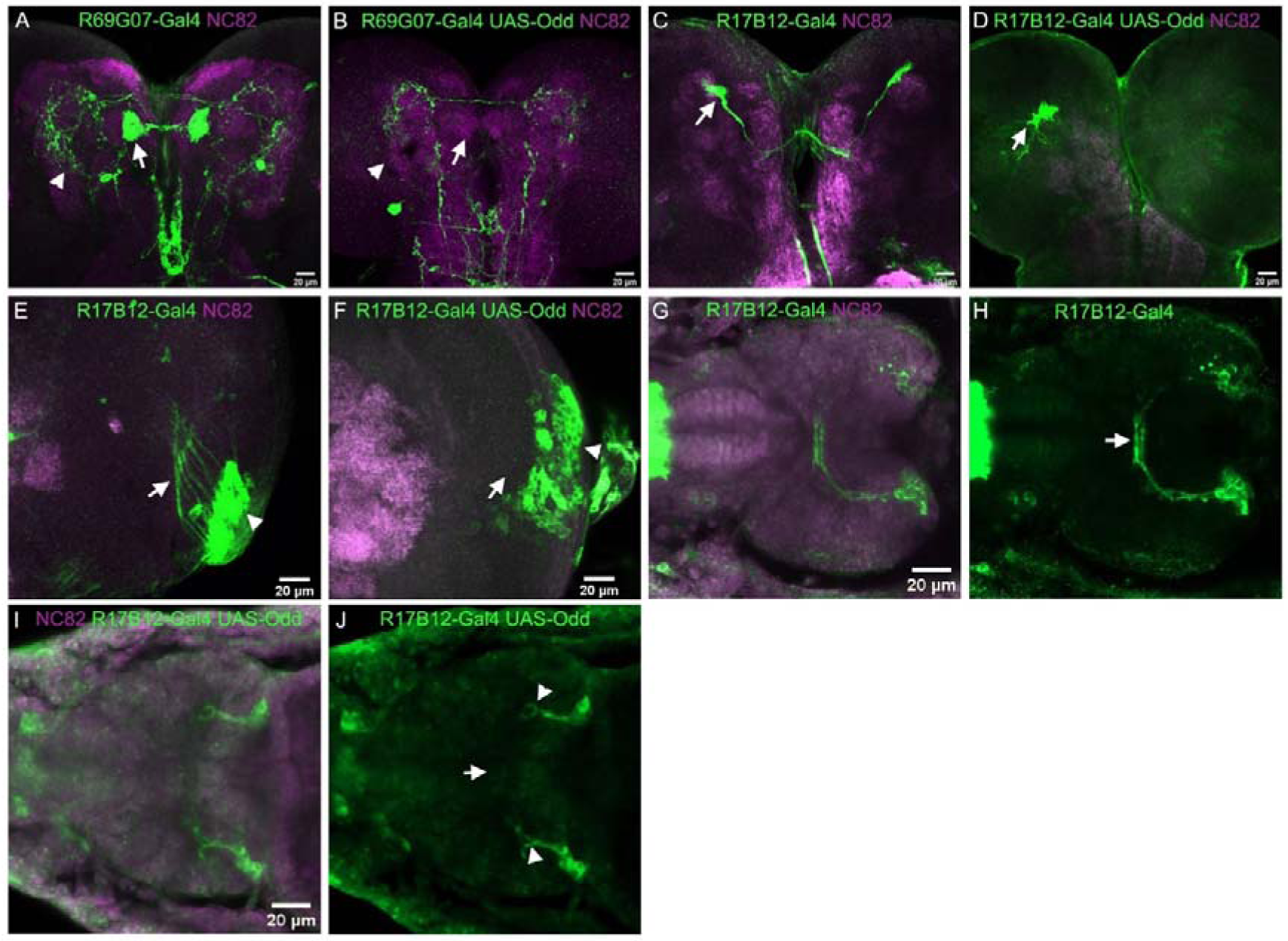
Miss-expressing Odd affects neurite growth and cell survival. Dorsal view of the brain. A-F) Anterior at the top, posterior at the bottom. G-J) Anterior to the left. A) The R69G07 Gal4 line drives expression in two neural populations. One which project anteriorly and across the midline arborizing in several compartments in the lateral part of the brain (arrowhead) and 1 which projects the medial part of the MB lobes (arrow). B) Projections to the MB lobe are missing in brains in which Odd is expressed using the R69G07 Gal4 line (arrow) whereas the neurites projecting anteriorly and ventrally maintain midline crossing although the ventral arbour is reduced in size (arrow). C) The R17B12 Gal4 line drives expression in a group of neurons situated anteriorly in the brain which project posteriorly before crossing the midline (arrow). D) These neurons are often missing in brains expressing Odd but sometimes survive in which case neurites appear to emerge randomly from the neurons and are severely stunted (arrow). E) The 17B12 Gal4 line is also expressed in optic lobe neurons (arrowhead) which project towards the laminar (arrow). F) Projections from these neurons are missing in brains miss-expressing Odd (arrow) and the neurons appear disorganised (arrowhead). G-H). The 17B12 Gal4 line is also expressed in the embryo in a neural cluster situated in the posterior part of the brain which project anteriorly and cross the midline (arrow). G) Both channels showing co-labelling with GFP antibody and NC82. H) Single channel showing GFP expression. I-J). Embryo expressing Odd using the 17B12 Gal4 driver. Expression is present in the embryo demonstrating that the neurons labelled by this line are born and form neurites when Odd is expressed. However, neurite projections are stunted (arrowheads) and do not cross the midline (arrow). I) Both channels showing co-labelling with GFP antibody and NC82. J) Single channel showing GFP expression.

### Increased Odd protein levels correlates with retention of synaptically targeted proteins to the cell body

During the morphological analysis of Odd over expression we noticed that Bruchpilot (Brp) antibody staining were noticeably stronger in the cell bodies of neurons that express RNAi or UAS-Odd compared to neurons that only express GFP in the control brains (Fig 7A-I) Brp localises to presynaptic terminals (Wagh, Rasse et al. 2006) and is used in this study and by many others to label the neuropil (e.g. Hartenstein, 2020 #32). In control brains (Odd-Gal4) as in wildtype, Brp protein is almost absent from the cell body (Fig 7A, B) whereas Odd neurons expressing RNAi 34328, RNAi VDRC and UAS-Odd show higher levels of Brp antibody staining in the cell bodies (Fig 7C-H). We quantified this by measuring the Brp antibody fluorescence intensity in the Odd expressing neurons and compared that to the fluorescence intensity immediately adjacent to the Odd neurons (Fig 7G, H). Brp antibody fluorescence intensity is similar in Odd neurons compared to adjacent tissue in control brains expressing GFP driven by Odd-Gal4 (Fig 7I). Brp antibody fluorescence level is significantly higher in Odd neurons expressing RNAi 34328, RNAi VDRC and Odd (Fig 7I). We next wondered whether this was unique to the presynaptic Brp or whether post synaptic associated proteins show a similar distribution. In order to address this question, we carried out immunofluorescence staining for Syndapin which localises to the post-synaptic membrane system (Kumar, Fricke et al. 2009). Similar to Brp, Syndapin is absent from the cell body in control brains expressing GFP in the Odd neurons (7J, K) but antibody fluorescence intensity is increased in Odd neurons expressing RNAi 34328, RNAi VDRC and Odd (Fig 7L-R). Quantifying fluorescence intensity levels between Odd neurons and adjacent tissue show that the Syndapin antibody fluorescence levels is significantly higher in Odd neurons expressing RNAi 34328, RNAi VDRC and Odd compared to control brains (Fig 7S). This data suggests that both Brp and Syndapin expression is increased in Odd neurons which over express Odd or alternatively is retained and not transported to pre and post synaptic sites.

**Figure 7.**
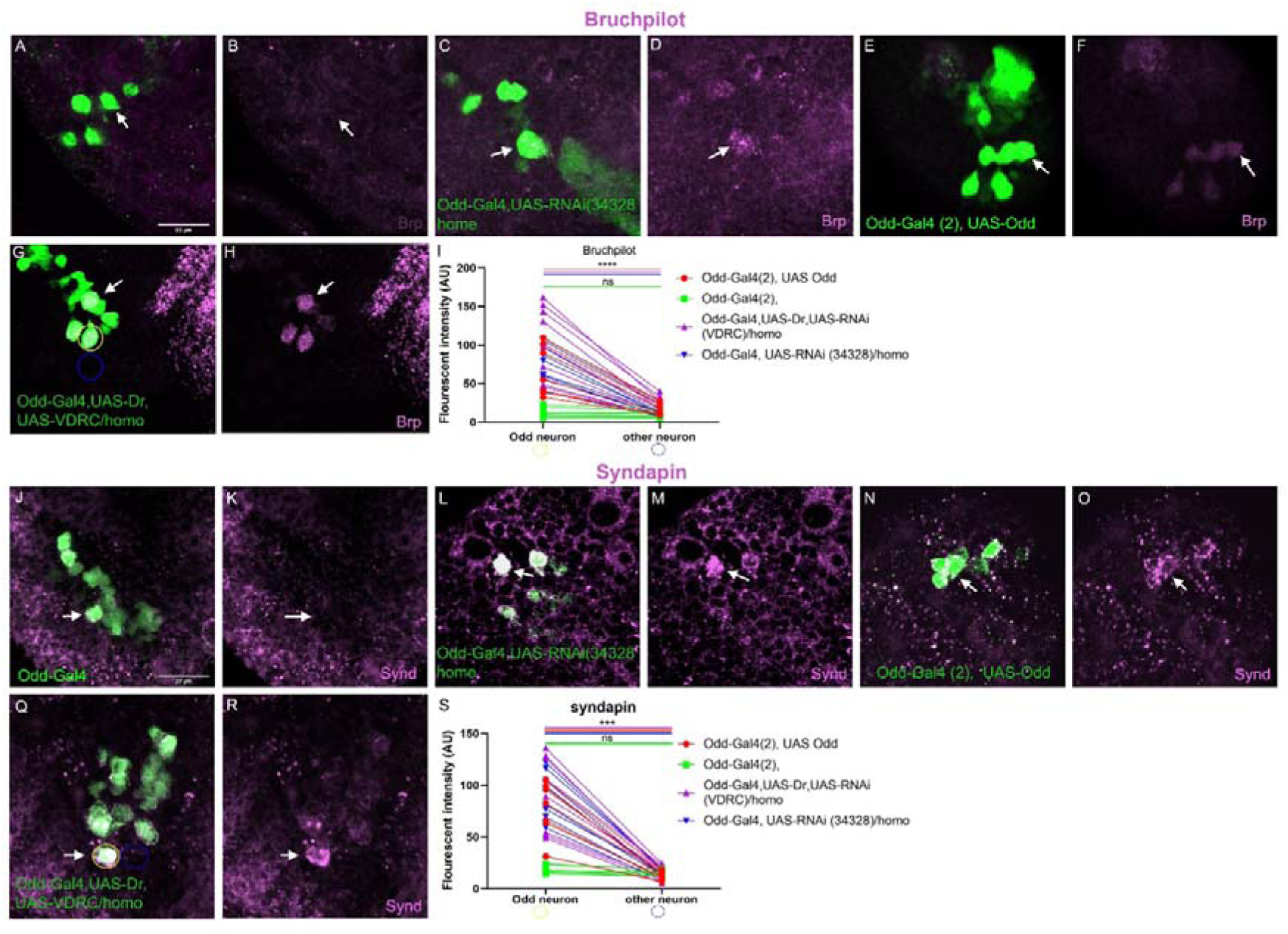
Increased Odd protein levels correlates with increased Brp and Syndapin immunofluorescence in the Odd neural cell bodies. Dorsal view of the brain. Anterior at the top, posterior at the bottom. A-H) Localisation of Brp (magenta) in brains. A,C,E,G) Both channels showing co-labelling with GFP antibody and NC82 (Brp). B,D,F,H) Single channel showing BrP expression. A-B) Control brains (Odd-Gal4) do not show specific localisation of Brp in the Odd neural cell body (arrow). Brp localises to the Odd neural cell bodies in C,D) brains expressing homozygous RNAi 34328, E,F) brains expressing 2 copies of UAS-Odd, and G,H) brains expressing homozygous RNAi VDRC (arrows). I) Quantification of Brp immunofluorescence in Odd cell bodies (marked by a yellow circle in G) and in an adjacent non-Odd expressing cell (marked by a blue circle in G). Immunofluorescence intensity is similar in Odd neurons and adjacent neurons in control brains whereas immunofluorescence intensity is significantly higher in Odd neurons compared to adjacent neurons in brains expressing 2 copies of UAS-Odd (red), homozygous expression of RNAi VDRC (purple) and homozygous expression of RNAi 34328 (blue) (2-way ANOVA). J-R) Localisation of Syndapin (magenta) in brains. A,C,E,G) Both channels showing co-labelling with GFP antibody and Syndapin. B,D,F,H) Single channel showing Syndapin expression. J-K) Control brains (Odd-Gal4) do not show specific localisation of Syndapin in the Odd neural cell body (arrow). Syndapin localises to the Odd neural cell bodies in L,M) brains expressing homozygous RNAi 34328, N,O) brains expressing 2 copies of UAS-Odd, and P,Q) brains expressing homozygous RNAi VDRC (arrows). R) Quantification of Syndapin immunofluorescence in Odd cell bodies (marked by a yellow circle in Q) and in an adjacent non-Odd expressing cell (marked by a blue circle in Q). Immunofluorescence intensity is similar in Odd neurons and adjacent neurons in control brains (Hao, Green et al.) whereas immunofluorescence intensity is significantly higher in Odd neurons compared to adjacent neurons in brains expressing 2 copies of UAS-Odd (red), homozygous expression of RNAi VDRC (purple) and homozygous expression of RNAi 34328 (blue) (2-way ANOVA)

**Figure 8.**
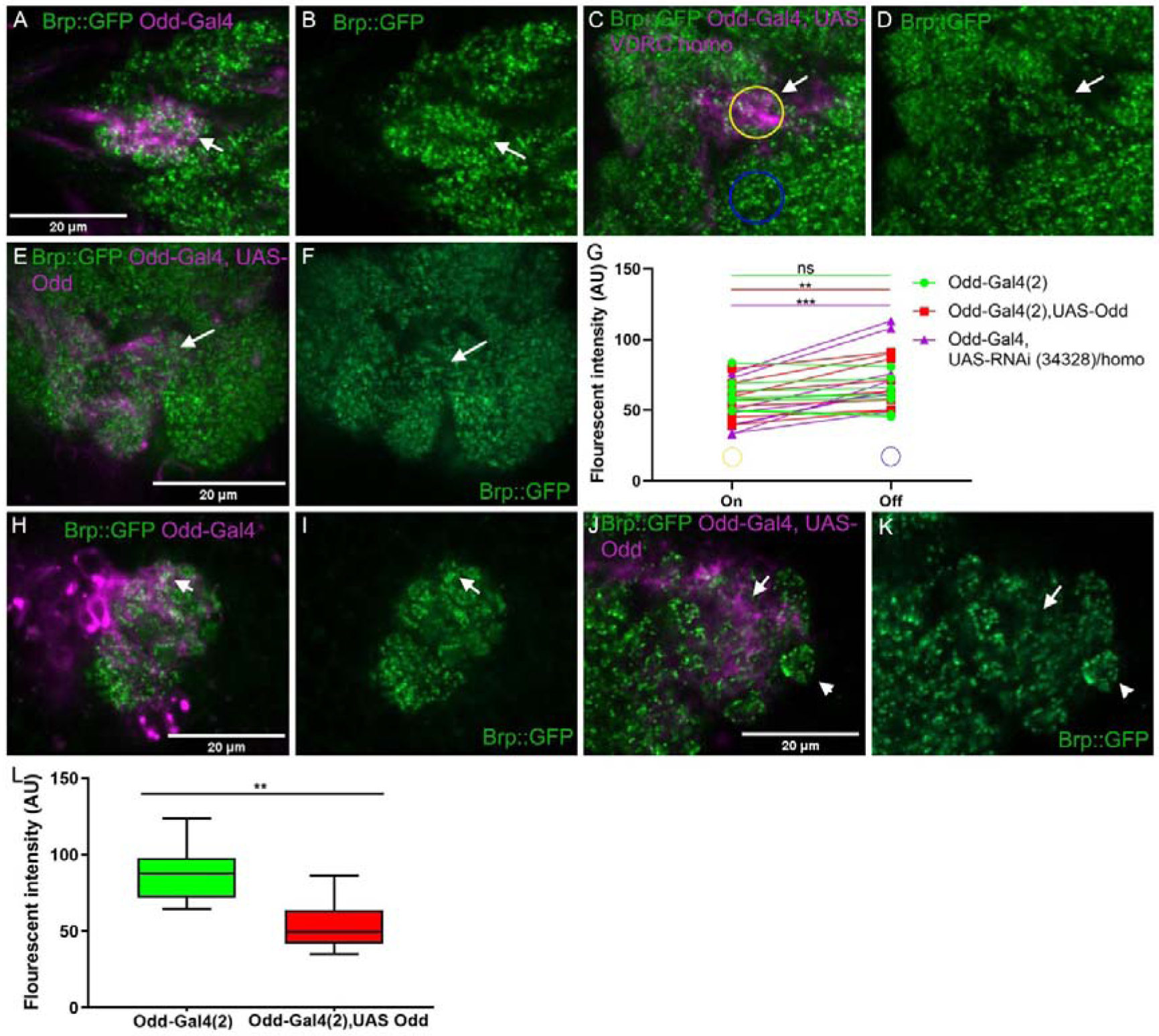
Increasing Odd protein levels is associated with a decrease in Brp::GFP at the axonal arbour. Dorsal view of the brain, anterior at the top, posterior at the bottom. A,C,E,H,J) Both channels showing co-labelling with RFP antibody (Odd neurite arbours) and Brp::GFP). B,D,F,I,K) Single channel showing Brp::GFP localisation. A-B) A-B) Brp::GFP localisation to the Odd axonal arbour in control brains (Odd-Gal4). There is no difference in GFP fluorescence levels between the arbour and adjacent tissue (arrows). Over-expression of Odd using the RNAi 34328 (C,D) or 2 copies of UAS-Odd (E,F) results in a reduction in GFP fluorescence in the axonal arbour compared to adjacent tissue (arrows). G) Quantification of GFP fluorescence in the Odd axonal arbour (On) compared to adjacent tissue (Off) shows a statistically significant decrease in GFP fluorescence in the axonal arbour (2 way-ANOVA). H-I) Brp::GFP fluorescence levels in the calyx of the control brains (Odd-Gal4) (arrow) is higher than that off GFP fluorescence in brains expressing 2 copies of UAS-Odd (J,K) (arrows). Areas of the calyx not innervated by the Odd dendrites show stronger GFP fluorescence compared to areas innervated by Odd dendrites (arrowhead).

**Figure 9.**
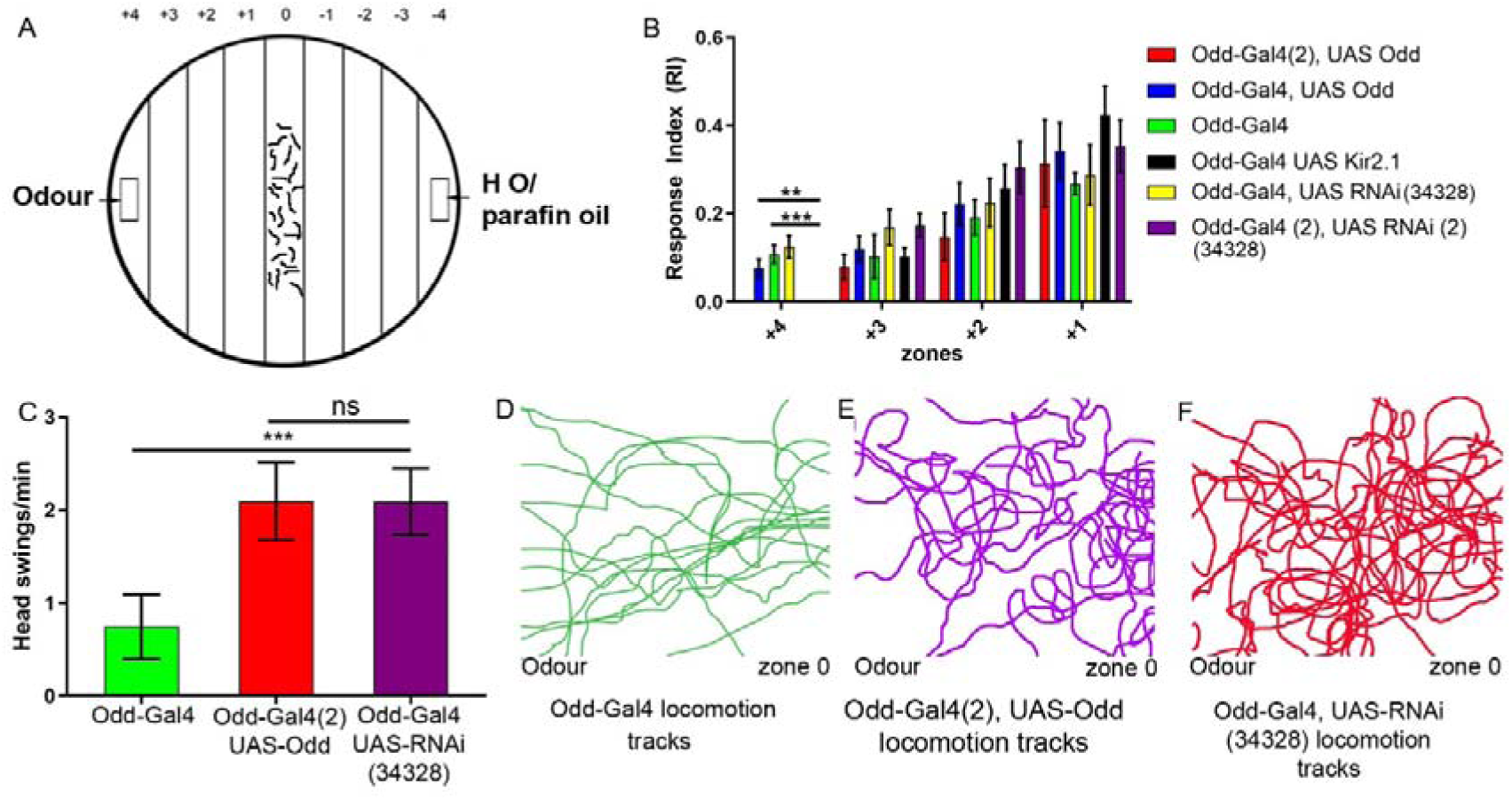
Over-expression of Odd renders Odd neurons non-functional. A) Schematic representation of chemotaxis arena. Odour is added to zone +4 and H_2_O added to the ࢤ4 zone. Larvae are placed in the 0 zone and allowed to wander for 5 min after which the number of larvae are counted in each zone. The proportion of larvae found in the 4 zones closest to the odour is calculated as number of larvae in + zone – number of larvae in -zone divided by total number of larvae and expressed as an RI index. A positive PI index indicates attraction to the odour whereas a negative RI index indicates repulsion. B) Bar graph showing the proportion of larvae found in each + zone. Larvae expressing 2 copies of UAS-Odd (red) and larvae expressing homozygous RNAi 34328 (purple) as well as Odd silenced larvae (black) do not reach the +4 zone and there are significantly more Odd-gal4 (Hao, Green et al.), Odd-Gal UAS-Odd (blue) and heterozygous expressing RNAi 34328 in the +4 zone (2 way-ANOVA).C) Larvae which express 2 copies of Odd and homozygous expressing RNAi 34328 carry out significantly more head swings/min compared to control larvae (Odd-Gal4) (1 way-Anova). D-F) Representation of locomotion tracks during chemotaxis form Odd-Gal, Odd-gal4 UAS-Odd (2) and homozygous RNAi 34328 respectively. Locomotion tracks are more convoluted when larvae express 2 copies of UAS-Odd and homozygous expression of RNAi 34328 compared to control larvae (Odd-Gal4)

### Increased *Odd-skipped* protein levels correlates with a decrease in Bruchpilot localisation in the axons

In order to address the effect of Odd over-expression on Brp localisation we chose a protein trap approach (Nagarkar-Jaiswal, Lee et al. 2015) in non-fixed tissue (see Materials and Methods). The advantage of this approach is that Brp, tagged with GFP, is driven by its endogenous promoter avoiding issues with miss localisation generated by UAS expression. Furthermore, despite the fact that we were careful to avoid bleed through by imaging each fluorophore separately we were still worried that using antibodies did not faithfully replicate actual protein levels. Since Brp localises to pre-synaptic terminals we focussed on the axonal arbour and measured GFP fluorescence co-localised with RFP driven by Odd-Gal4. And compared GFP fluorescence intensity in the arbour to that of GFP fluorescence in adjacent tissue in the same sample. There is no visible difference in GFP fluorescence intensity between the Odd neural arbour and adjacent tissue in control brains (Odd-Gal4) (Fig 8A, B). However, both homozygous expression of RNAi (34328) (Fig 8C,D) and expressing 2 copies of UAS-Odd in the Odd (Fig 8E,F) neurons significantly reduces the number of Brp::GFP puncta and therefore GFP fluorescence levels in the axonal arbour. Comparing fluorescence intensity levels between Odd arbours (ON) and adjacent tissue (OFF) show that there is a significant decrease in GFP florescence level at the Odd neural arbour (Fig 8G). There are some Brp::GFP puncta localised to the axonal arbour suggesting that Odd over expression does not prevent transport to the synapse. The reduction in Brp::GFP puncta could be a result of reduced synapse formation in Odd neurons potentially because the axons have not arborized in the correct part of the brain which could prevent synapse formation between Odd neurons and their postsynaptic target. In order to address this, we focussed on the dendritic arbour in the calyx. The dendritic arbours in the calyx are missing in brains express RNAi in the Odd neurons but over-expression of Odd causes only a partial loss of dendritic arborisation in the calyx. Importantly this phenotype does exhibit some correctly localised dendrites which should allow for synapse formation between Odd neurons and their presynaptic targets. We therefore measure Brp:: GFP fluorescence intensity in the entire calyx and compared calic GFP levels between control brains and that of brains expressing 2 copies of UAS-Odd in the Odd neurons (Fig 8H-L). Since Brp::GFP localises to all presynaptic sites there will still be considerable Bruchpilot::GFP fluorescence in the calyx since we predict that other synapses such as the synaptic connections between PNs and Kenyon cells are normal in brains in which Odd is over-expressed in the Odd neurons. Despite this we found a significant reduction in calic GFP fluorescence in brains over-expressing Odd (Fig 8J,K) compared to control (Odd-Gal4) brains (Fig 8H,I). Interestingly this reduction seems to be localised specifically to areas surrounding the Odd dendritic arbours whereas micro glomeruli not innervated by Odd dendrites have more GFP puncta (Fig 8J, K). This data suggests that manipulating Odds-kipped protein levels affects synapse formation.

### Increasing *Odd-skipped* protein levels disrupts Odd neural function

Increasing Odd protein levels either by RNAi expression or over expression of Odd clearly affects Odd neurite morphology and synapse formation and we therefore addressed whether these changes would affect the function of Odd neurons. We have previously shown that silencing Odd neurons (Odd silenced larvae) impair chemotaxis by reducing odour concentration discrimination (Slater, Levy et al. 2015). Chemotaxis is the ability of most animals to detect the orientation and magnitude of an odour gradient to direct navigation towards an odour source. *Drosophila* larvae adopt stereotyped behaviours during chemotaxis including lateral head swings which are thought to allow the larvae to sample the odour space and measure the direction of the odour gradient (Gomez-Marin, Stephens et al. 2011). If larvae are moving away or perpendicular to the odour source lateral head swings will frequently be followed by a directional turn which is more likely to reorient the larvae towards the odour source than away from the odour source. Larvae in which Odd neurons are silenced are worse at localising an odour source compared to control larvae. Frequency of lateral head swings and turn rate are significantly increased in larvae in which the Odd neurons are silenced compared to control larvae resulting in more convoluted navigation tracks. Based on this data we reasoned that if morphological changes affect neural function, we expect to observe a similar behavioural phenotype to that of larvae in which we have silenced the Odd neurons. We used the larval chemotaxis assay previously described (Fig 9A) (Slater, Levy et al. 2015) and filmed larval behaviour during chemotaxis. The chemotaxis arena is divided into 9 zones; 4 zones on the odour side of the arena and 4 zones on the non-odour side of the arena separated by a 0 zone in which the larvae are placed. Larvae are allowed to navigate the odour gradient for 5 min after which the number of larvae are counted in each zone. The proportion of larvae in each odour zone is expressed as a response index (see materials and methods). This design allows us to measure how well larvae can navigate the odour gradient by precisely quantifying their position in relation to the odour source at the end of the experiment. Larvae in which we silence Odd neurons do not reach the odour source in zone +4 but do navigate towards the odour source represented by a positive RI index in zone +3, +2, +1 (Fig 9B). Similar to silencing Odd neurons, larvae that are homozygous for RNAi (34328) and larvae containing 2 copies of UAS-Odd do not reach the odour source in zone +4 but do navigate towards the odour. Odd-gal4 (control) larvae and larvae that express RNAi (34328) heterozygous (Odd-Gal4 UAS-RNAi (34328) can reach the odour source and have a similar RI index to that of control larvae (Odd-Gal4) A significantly smaller proportion of Odd-gal4 UAS-Odd larvae can also reach the odour source. This would suggest that increasing Odd protein levels affect Odd neural function. To further confirm this result, we asked whether homozygous RNAi (34328) and Odd-Gal4 (2) UAS-Odd larvae show similar changes in behaviour during chemotaxis as that previously observed in Odd-silenced larvae. Similar to Odd silenced larvae, Odd-Gal4, UAS-Odd (2) and homozygous RNAi (34328) larvae carry out significantly more head swings/min compared to control larvae (Odd-Gal4) (Fig 9C) and their navigation tracks are more convoluted compared to control larvae (Fig 9D-F). This data shows that increasing Odd protein levels has a similar effect on neural function as silencing Odd neural activity suggesting that the morphological changes coursed by elevated Odd protein levels renders the Odd neurons non-functional.

## Discussion

### Odd-skipped plays an important role in neurite morphology

Our data show that manipulating Odd protein levels affect neurite morphology and this effect appears to be dependent of neural sub-population. Increasing Odd protein levels in Odd neurons both through RNAi expression and over expression of Odd abolish dendritic arbour formation and axonal arbours are miss localised and appear defasciculated. The axons that cross the midline are missing or greatly reduced. Wildtype Odd neurons are tightly fasciculated and the axonal arbour in the CPM is very contained. As such this phenotype does resemble that of de-fasciculation suggesting that Odd could affect the expression of fasciculation proteins. We tested the expression using antibody staining of NCAD, FassII, Connectin and Semaphorin 1a which have all been associated with fasciculation in *Drosophila* (Yu, Huang et al. 2000, Spindler, Ortiz et al. 2009). We did not observe any obvious decrease or increase in the expression levels of these molecules in Odd neurons when Odd protein levels were increased. However, we cannot exclude the possibility that Odd may affect some other yet to be identified protein involved in fasciculation in *Drosophila.* On the other hand, our data show that synapse numbers have decreased in Odd neurons in which Odd protein levels are higher. Interestingly this is associated with retention of two synoptically targeted proteins; Bruchpilot and Syndapin to the cell body. This phenotype could be due to either a failure in the transport of these proteins to the axonal terminal or alternatively that the Odd neurons cannot make synaptic contact with the appropriate post-synaptic targets and synaptically targeted proteins therefore build up in the cell body. Although we did not investigate axonal and dendritic transport in detail, we can confirm that Odd neurons in which we have increased Odd protein levels contain actin bundles that have a similar morphology to that of neighbouring neurons. Instead we favour the latter hypothesis which is supported by our behavioural data. As axonal arbours are miss-localised and bigger it is likely that they cannot make synaptic contact with the correct post synaptic targets. Our behavioural data demonstrates that increasing Odd protein levels have a similar behavioural effect to that of silencing Odd neurons strongly suggesting that increasing Odd protein levels abolish neural function. That would certainly be the case if Odd neurons are not connected to the appropriate post synaptic neurons.

### RNAi against Odd upregulates Odd protein levels

RNAi are traditionally used to knock down protein levels and it was surprising to us that both RNA construct showed increase in Odd protein levels. This does highlight the importance of verifying protein levels when using RNAi for knock down purposes. How can RNAi upregulate protein levels? Odd can act as a transcriptional activator or repressor when bound to Groucho (Goldstein, Cook et al. 2005). Groucho is ubiquitously expressed in the CNS (Knust, Tietze et al. 1987) and it is therefore possible that Odd bound to Groucho predominantly act as a transcriptional repressor in the CNS. We hypothesis that Odd either directly or indirectly regulate its own transcription but not to turn expression on or off but rather to strictly maintain Odd protein levels. RNAi through an unknown mechanism disrupts the repression of Odd-skipped in place to prevent high levels of Odd expression. Removing Odd mRNA through RNAi lifts the repression of Odd-skipped expression allowing for an increase in Odd transcription. We speculate that this causes cyclic fluctuation in Odd-protein expression which may explain why protein levels as measured using western blots varies between samples. Since we used 3 different RNAi construct that all produced similar phenotype we think it is unlikely that the increase in Odd protein levels are due to some off-target effect of the RNAi since it would be unlikely that all 3 RNAi constructs have similar off-target effects. Interestingly loss-of Odd in Odd neurons has a more severe phenotype than increasing Odd protein levels where neurons either fail to produce neurites or neurites are small and stunted with no branching. This is also seen in some neural sub populations. For example, the optic lobe neurons labelled by the 17B12-Gal4 line completely fail to form axonal projection into the lamina. Thus, it appears that both loss of Odd function and upregulation of Odd course neurite defects further suggesting that precise levels of Odd protein is important for neural growth.

### Manipulating Odd expression can cause neural death

Our data suggests that manipulating Odd protein levels can cause cell death both in loss-of-Odd function and miss expression. This observation is supported by our MARCM study where we never see MARCM Odd null full lineage clones suggesting that induction of Odd null clones in the NB results in either cell death or cell cycle arrest. If the Odd null NB clone was viable, we would expect to observe Odd mutant descendants of the NB which we never do. Rather we consistently see 1 or 2 neurons which are homozygous for the null allele suggesting that induction of Odd null clone is viable in the Mother Ganglion Cell. The miss expression phenotypes also suggest that manipulating Odd protein levels can cause cell death. This is based on the observation that in many of the lines we used to miss express Odd one or more neurons are missing at 3^rd^ instar larvae. The Janelia farm collection of Gal4 driver lines each contain a 1 Kb fragment of an enhancer and it is possible that Odd could be regulating expression of the gene whose enhancer drives the Gal4 line in which we observe missing neurons. If Odd skipped binds to the enhancer region to repress expression, we would expect GFP expression to be down regulated giving the impression that the neuron is missing. However, we used several different Gal4 lines and in the majority of these lines, neurons were missing. It is unlikely that Odd should control expression in all the lines used for the miss-expression study. In 3 of the Gal4 lines we analysed, we identified expression of the driver line in embryos and in all 3 lines the neurons were present at embryonic stages. This data demonstrates that the neurons are born and begin to project neurites but likely die at some point in larval life. Interestingly, expressing Odd in all postmitotic neurons using the Elav-Gal4 driver line is embryonic lethal. Unlike the Odd-Gal4 line and some of the Janelia Farm lines Elav-Gal4 is only expressed in the CNS and we can therefore conclude that the lethality is due to expression of Odd in neurons. Although neurites are present in embryos expressing Odd the number of midline crossing fascicles are greatly reduced supporting the observation that some neural sub types are not affected by Odd expression. However, it is likely that if Odd disrupts neurite projections from a large population of neurons this would cause lethality especially since motor neurons would also be affected by expressing Odd using the ELAV driver.

In this study we have uncovered a novel role of Odd in the control of neurite growth and morphology. Our data supports the finding from vertebrate cancer studies that over expression of Odd correlates with cell cycle arrest and lack of proliferation. Thus, the role of Odd in proliferation is conserved demonstrating that the role of Odd in cell cycle arrest and proliferation can be studied in flies.

